# A like-for-like comparison of lightweight-mapping pipelines for single-cell RNA-seq data pre-processing

**DOI:** 10.1101/2021.02.10.430656

**Authors:** Mohsen Zakeri, Avi Srivastava, Hirak Sarkar, Rob Patro

## Abstract

Recently, Booeshaghi and Pachter (1) published a benchmark comparing the kallisto-bustools pipeline (2) for single-cell data pre-processing to the alevin-fry pipeline (3). Their benchmarking adopted drastically dissimilar configurations for these two tools, and overlooked the time- and space-frugal configurations of alevin-fry previously benchmarked by Sarkar et al. (3). In this manuscript, we provide a small set of modifications to the benchmarking scripts of Booeshaghi and Pachter that are necessary to perform a like-for-like comparison between kallisto-bustools and alevin-fry. We also address some misuses of the alevin-fry commands and include important data on the exact reference transcriptomes used for processing^1^. Using the same benchmarking scripts of Booeshaghi and Pachter (1), we demonstrate that, when configured to match the computational complexity of kallisto-bustools as closely as possible, alevin-fry processes data faster (~2.08 times as fast on average) and uses less peak memory (~ 0.34 times as much on average) compared to kallisto-bustools, while producing results that are similar when assessed in the manner done by Booeshaghi and Pachter (1). This is a notable inversion of the performance characteristics presented in the previous benchmark.

## Introduction

Alevin-fry (3) is a new pipeline for single-cell RNA-seq pre-processing, which is currently being developed. While there are many relevant design decisions and performance implications we hope to discuss in detail in the preprint describing alevin-fry, one crucial aspect motivating the development of the alevin-fry pipeline is to allow testing the effect of different algorithmic choices on the gene expression estimates eventually produced by the pipeline. For example, alevin-fry exposes both a selective-alignment (4, 5) mode and pseudoalignment (6) with structural constraints mode for mapping reads. Further, after read mapping, the alevin-fry tool exposes multiple algorithms for generating a permit list (sometimes called a “whitelist”) of barcodes corresponding to what are believed to be high-confidence cells, and for resolving UMIs into counts. When applying any of the probabilistic methods it implements for UMI resolution, alevin-fry also allows assessing quantification uncertainty in the estimated counts via a bootstrapping procedure that can output either the bootstrap samples, or their summary statistics.

Exploring these different algorithms in a unified framework is an important task to optimize the pre-processing of single-cell sequencing data, and there may not be a single algorithm that is best suited to all different single-cell technologies. For example, while the benefits of selective-alignment and the use of an expanded index in the processing of bulk RNA-seq data have been highlighted in a growing number of scenarios (4, 7, 8), these tradeoffs have not been thoroughly explored in the context of single-cell (and particularly tagged-end) data. Given that the majority of common tagged-end single-cell analyses are performed at the gene rather than transcript level, and in light of the extensive use of techniques like unique molecular identifier (UMI) tagging, it may be the case that different tradeoffs in mapping specificity versus speed are appropriate or desirable — indeed, an argument for simpler but faster methods in this space has been made by Melsted et al. (2) and in subsequent work by the same authors. Likewise, the effect of different approaches for UMI error correction and UMI resolution (and how they may interact with different read mapping strategies) has not been thoroughly evaluated across many different single-cell technologies, to understand if, and when, different approaches may lead to different results in downstream analysis.

In the alevin-fry poster (3), we described the results of benchmarking STARsolo (9), kallisto-bustools (2) and alevin-fry (3), running the latter tool with a number of different configurations of read mapping algorithm and UMI resolution algorithm. We observed that the “fast” configurations of alevin-fry tested in (3), which adopt some of the major simplifications argued for by Melsted et al. (2), are faster than kallisto-bustools, and that all of the configurations tested there use less peak memory. The recent preprint of Booeshaghi and Pachter (1) omits all of the fast and memory-frugal configurations tested in Sarkar et al. (3), and instead compares the time and memory requirements of only the most computationally- and memory-intensive configuration of the alevin-fry pipeline to the kallisto-bustools pipeline.

We are encouraged that others in the community are eager to try out new tools like alevin-fry for the pre-processing of single-cell data, and we recognize that fairly comparing new pipelines to existing ones can be a difficult task in the absence of sufficient documentation and tutorials. Admittedly, we have not yet produced sufficient tutorials or documentation for alevin-fry given that our efforts have been in continuing to develop the tool itself. At the same time, it is not possible to “faithfully” follow recommended practice (1) when the best practices have not yet been established for a fledgling method; in such a case, benchmarking multiple configurations (especially those that have already been tested in previous benchmarks (3)) may be a reasonable approach. Spurred by Booeshaghi and Pachter (1), we have now created a simple-to-follow tutorial for speed-optimized single-cell pre-processing using alevin-fry (https://combine-lab.github.io/alevin-fry-tutorials/2021/running-alevin-fry-fast/). Here, we benchmark this workflow for alevin-fry (using the same versions of salmon (1.4.0) and alevin-fry (0.1.0)^2^, and kallisto(0.46.2)-bustools(0.40.0) adopted in (1)).

## Methods

In order to assess how the results of the benchmark proposed by Booeshaghi and Pachter (1) change when a like-for-like comparison between alevin-fry and kallisto-bustools is carried out, we start with the experimental framework introduced in that preprint and describe here the necessary modifications to the benchmarking scripts that were made.

The first difference we note is the versions of the references used for quantification. The repository provided by Booeshaghi and Pachter (1) (https://github.com/pachterlab/BP_2021/ commit `d0549e0351c4875428b153c7804f21eed7fa82eb`, which was the version available when the preprint was published) lacked both the specific reference sequences used and the URLs from which these reference sequences were obtained^3^. The paper refers the reader to (2), wherein the relevant metadata is contained in an Excel file, which still lacks adequate specificity (*e.g*. it lists the *Caenorhabditis elegans* transcriptome used only as “modified ws260”). Thus, in this manuscript, we have adopted the following procedure for normalizing the reference transcriptomes. For human, mouse, and combined human/mouse data, we have used the latest reference bundles provided by 10x Genomics as of Jan 29, 2021 (named as “2020-A”), and extracted the transcriptomes from the provided genomes and respective GTF files using gffread (10). For all other organisms, we have adopted the latest Ensembl (11) reference transcriptomes for each organism. For *Danio rerio, C. elegans, Drosophila melanogaster* and *Rattus norvegicus* this is from release 102; for *Arabidopsis thaliana* it is from release 49.

These updated reference transcriptomes lead, in some cases, to quite different memory usages from those reported in (1). For the alevin-fry pipeline, this is largely explained by the fact that we index the *same* reference sequences as used for kallisto (6) (that is, we do not compare indexing the transcriptome in kallisto to indexing the transcriptome and genome in salmon (12)). However, the increased memory usage of kallisto-bustools likely stems from variation in the specific reference transcriptomes used. For example, using the current 10x reference transcriptome for GRCh38 (2020-A, from https://cf.10xgenomics.com/supp/cell-exp/refdata-gex-GRCh38-2020-A.tar.gz), the peak memory usage of kallisto becomes ~7GB during mapping (Fig. 1), rather than the ~4GB reported in (1) (while the peak memory usage of salmon during mapping reaches ~ 1.7GB). Our version of the repository contains a file called `gather_refs.sh` with the commands used to obtain these reference transcriptomes.

**Fig. 1.**
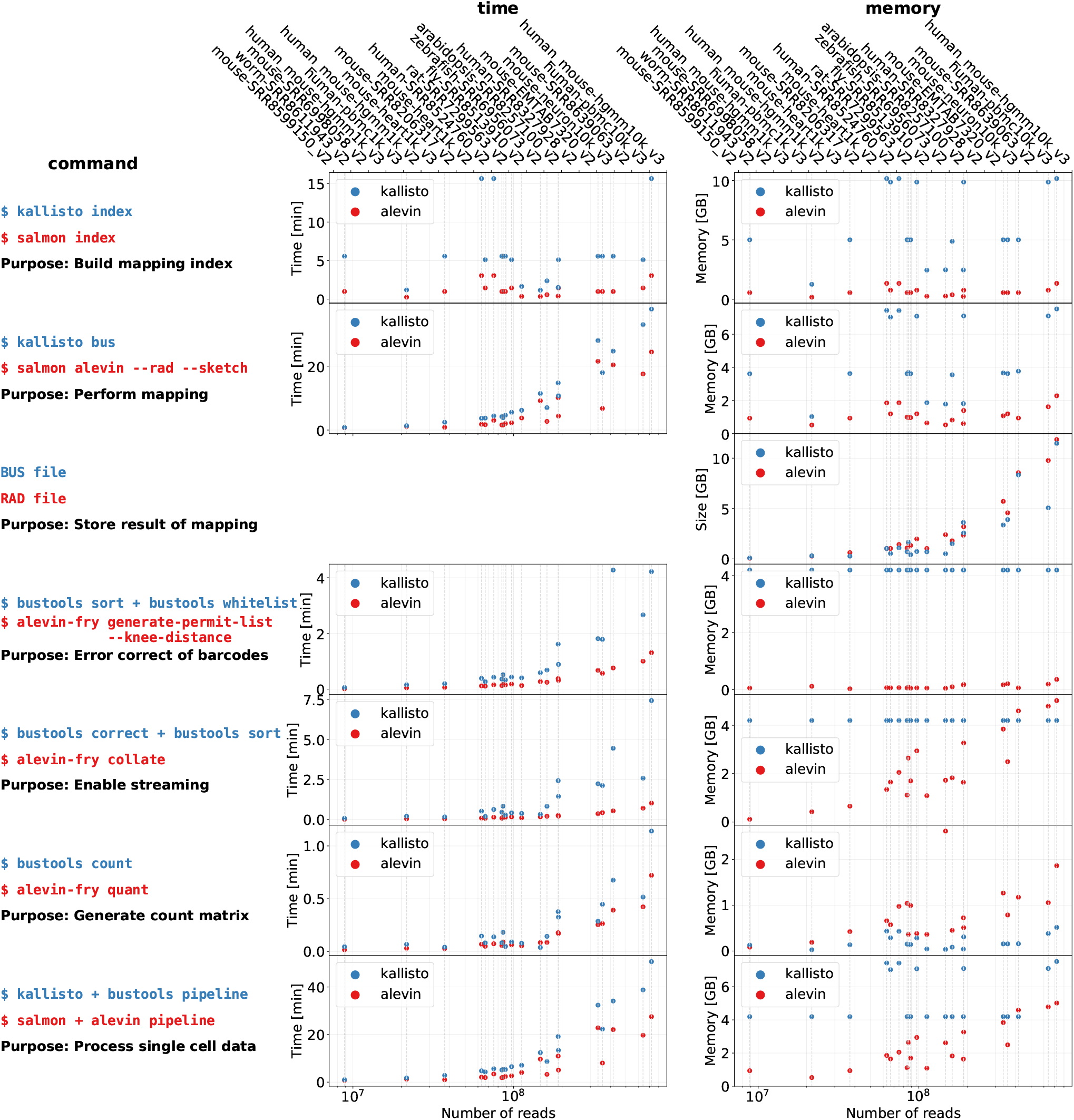
The time and memory used by the relevant steps of the alevin-fry and kallisto-bustools pipelines for pre-processing the 20 diverse tagged-end single-cell RNA-seq datasets used in (1). The plots are generated using the `analysis/notebooks/memtime.ipynb` notebook.

Furthermore, the following additional modifications have been made to the benchmarking, which otherwise remains the same as was performed in (1). We run alevin with the `--sketch` flag when producing the mapping file (called a RAD file); this uses pseudoalignment (6) with structural constraints rather than selective-alignment^4^. We do not consider a configuration of either kallisto-bustools or alevin-fry that corrects to or uses the full 10x permit list. In the original benchmark, the authors used alevin-fry with the `-b` flag, treating the full list of 10x barcodes as a filtered permit list; however the `-b` flag is meant to accept a list of barcodes corresponding to high-confidence cells that have passed external filtering. Passing the full 10x barcode list to the `-b` flag is neither intended nor currently supported in alevin-fry (though we are planning to add this functionality), which we have now clarified in the documentation — as we had previously clarified this same point in the alevin (13) documentation. We have both methods generate their own permit list, and perform quantification on their corresponding filtered cells. We pass the `-d fw` flag to alevin-fry’s `generate-permit-list` step rather than `-d either`, as the RAD file records the orientation of each read with respect to the target transcript, and all the technologies evaluated here expect the second read to map to the transcriptome in the forward orientation. Mappings in an unexpected orientation should be filtered. We have used the `cr-like` resolution strategy when invoking the alevin-fry `quant` command; this implements a simple but fast UMI resolution algorithm that breaks ties by UMI frequency alone and discards reads for which a most frequent unique gene cannot be determined.

We have also removed the step of the pipeline that converts the RAD (respectively BUS) file into a text format. The binary to text conversion may be useful for debugging purposes, but is not a standard or necessary part of these pre-processing pipelines, as the BUS and RAD files are primarily intended for the storage and processing of data rather than human inspection. Further, contrary to the supposition of Booeshaghi and Pachter (1), this conversion is likely a case where language choice, and usage of standard language idioms, leads to different performance characteristics. Unlike C++, Rust places the standard output stream behind a lock to ensure threadsafe access, a decision that imposes a cost for programs that are heavy on writing to the standard output stream in a line-oriented manner when standard idioms are used. While we do not view the optimization of the command that dumps a RAD file to text as particularly high-priority, we will nonetheless explore making use of unsafe C system calls in this command until a comparable solution is exposed natively in Rust.

The benchmarking scripts used to produce the results described here can be found at https://github.com/COMBINE-lab/BP_2021-lfl (these are the same as the benchmarking scripts of https://github.com/pachterlab/BP_2021 at commit `d0549e0351c4875428b153c7804f21eed7fa82eb` with the modifications described above). We encourage users to run these benchmarks for themselves, and welcome feedback and suggestions.

Despite the additions and modifications we describe here, neither our repository nor the original repository of Booeshaghi and Pachter (1) enable full reproducibility without non-trivial effort or investigation. One complication is that there existed multiple candidate scripts for performing specific steps of the data analysis within different directories of the repository, and none had complete or adequate instructions for generating the plots. For example, there exist multiple versions of the `run_gsea_bar_full.R` script for performing gene set enrichment analysis, which each required building certain sub-directories in the main directory of the repository in order to be executed without any errors. Eventually, we used the `run_gsea_bar_full.R` script located within `analysis/notebooks` rather than the one located in `analysis/scripts/code`, since the latter version had hard-coded paths and no central way to uniformly and globally change the working directory (*e.g*. https://github.com/pachterlab/BP_2021/blob/e87e98713bf7967d2fa22716dbbebd10609c1dd9/analysis/scripts/code/gsea_bar_full.R#L39). After providing the required data, we ran `mkdata.py` and `mk-plot.py` within the `analysis/scripts/code` directory to prepare the plots for comparing the gene count estimates provided by both tools. Furthermore, since we benchmarked an unmodified version of alevin-fry, we had to modify the `mkdata.py` script to load a single column file as alevin’s permit list (which we took from the `quants_mat_rows.txt` file accompanying each cell by gene count matrix), and also to remove the lines which were intended for dealing with decoy aware results. For producing the time and memory plots, we used the `memtime.ipynb` notebook located in `analysis/notebooks` after making the required modifications to compare the time and memory of the relevant steps in both tools. While we have addressed any issues as we encountered them, and have documented how we have run this pipeline, we have not undertaken the effort of fully removing all barriers to “trivial” reproducibility, as it is outside the scope of the current work.

Finally, we also note that the benchmark of Booeshaghi and Pachter (1) focuses only on comparing kallisto-bustools to a single configuration of alevin-fry, excluding other relevant tools like STARsolo (9), which is a fast, flexible, and popular tool for the pre-processing of tagged-end single-cell data. The benchmark also omits another recently-published, lightweight-mapping based tool, Raindrop (14), from the benchmark (though, seemingly, this would currently have to be restricted to 10x chromium v2 data). A more extensive benchmark, including other tools, is likely to provide greater value to the broader community. However, the primary focus in this manuscript is to highlight the effect on the original benchmark that results from running the tools considered therein in a like-for-like configuration. Thus, we have not added STARsolo or Raindrop to the current benchmark, though it may provide a useful perspective on these tools to the broader community.

All experiments were performed on a server with dual Intel Xeon CPUs (E5-2699 v4), each with 22 cores clocked at 2.20 GHz, 512 GB of 2.4GHz DDR4 memory, and an array of 8 3.6TB Toshiba MG03ACA4 HDDs configured as independent disks.

## Results

Fig. 1 shows the overall time and peak memory taken by both alevin-fry and kallisto-bustools when pre-processing the 20 diverse tagged-end single-cell 10x chromiumdatasets evaluated in (1). Alevin-fry is faster than kallisto-bustools on all datasets (between ~1.28 and ~ 2.87 times as fast, and ~2.08 times as fast on average). Also, alevin-fry uses less peak memory than kallisto-bustools on 19 of the 20 datasets tested, with the peak memory of kallisto-bustools ranging from ~91% of that used by alevin-fry to ~8 times that used by alevin-fry (kallisto-bustools used ~2.92 times as much peak memory on average).

In addition to the overall runtime and peak memory usage (bottom row of Fig. 1), the figure also shows the time and memory required for the main steps of the pipelines. While there is not a perfect correspondence between the specific set of commands used by the two tools, the fundamental steps include mapping the reads, generating a permit-list of valid filtered barcodes (with each method using its own algorithm to infer the filtered set of corrected barcodes), rearranging the mapping information for all records having the same corrected barcode so that they are adjacent in the resulting file, and applying a UMI resolution algorithm to obtain a gene-by-cell count matrix.

Looking across the datasets, some general characteristics emerge. If one evaluates the ratio of the total runtime of kallisto-bustools to the total runtime of alevin-fry, one observes that alevin-fry is faster in the processing of every dataset, with a speedup (*i.e*. runtime of kallisto-bustools/ runtime of alevin-fry ratio) ranging from ~1.28 up to ~2.87 (with an average runtime speedup of ~2.08). If one evaluates the same ratio in terms of peak memory usage instead of total runtime, a similar trend emerges. In 19 of the 20 datasets tested here, the kallisto-bustools pipeline exhibits a higher peak memory usage than alevin-fry. In the mouse-SRR8639063_v2 dataset, kallisto-bustools’ peak memory usage reached 91% of that of alevin-fry (with alevin-fry requiring a maximum of ~4.6GB^5^ of memory and kallisto-bustools requiring ~ 4.2GB of memory). In every other dataset, kallisto-bustools usedmore peak memory than alevin-fry, with the kallisto-bustools pipeline using at most ~8 times as much peak memory and, on average, ~2.92 times as much peak memory as alevin-fry. The peak memory usage of both tools reached their respective maxima on the hybrid human-mouse dataset, where the peak memory usage of kallisto-bustools is ~7.5GB (which occurs during pseudoalignment) and the peak memory usage of alevin-fry is ~5GB (which occurs during mapping record collation).

While the step of indexing only has to be done once per reference sequence (*i.e*. with each new organism, or when the reference annotation is updated), we also evaluate the time and memory required to build all indices used in these experiments. This is important, since the peak memory usage during indexing may dictate whether the index can be built on the same machine used for subsequent quantification, or if it must be constructed on a machine with more memory. Fig. 1 shows that, as with the pre-processing of reads, alevin-fry is faster and uses less memory for index construction for each reference considered. The slowest index construction for both tools was for the human-mouse combined transcriptome, where the kallisto-bustools pipeline took ~ 15.6 minutes and required ~ 10.2GB of memory, while indexing this transcriptome with the alevin-fry pipeline took ~3 minutes (a ~ 5.2 times speedup compared to kallisto) and ~ 1.3GB of memory (~13% of the the memory usage of kallisto). When evaluating the time differences, it is important to note that the alevin-fry pipeline can make use of multiple threads when indexing (here we used 10 as in (1)), while the indexing in the kallisto-bustools pipeline is currently restricted to a single thread. The memory usage in the alevin-fry pipeline does not vary considerably with the number of threads used during indexing. The peak memory reduction during indexing and mapping in alevin-fry arise primarily due to alevin-fry’s use of the pufferfish (15) index, while a number of different factors at both the implementation and design level contribute to the runtime improvements.

When assessing the same summary statistics and count comparisons considered by Booeshaghi and Pachter (1) to evaluate the similarity of the resulting quantifications, we find that the cell by gene count matrices produced by both tools are similar under these metrics (Fig. 2). As is expected, these evaluations show that the data summaries are more similar than in the configuration tested in (1). In that comparison, Booeshaghi and Pachter (1) claim that differences in resulting gene expressions between the configurations of the tools tested therein are “irrelevant for downstreamanalysis” (presumably implying all possible downstream analyses). It is not clear how these comparisons justify such a sweeping claim. Yet, while these comparisons do not necessarily imply that no differences will manifest in downstream processing of the alevin-fry quantified data compared to the kallisto-bustools quantified data, they do suggest that the differences that may arise under this configuration of alevin-fry are likely to be less extreme than differences that may arise in the configuration tested in (1). We also note that, while Booeshaghi and Pachter (1) observe no significant gene sets found when comparing the quantifications of kallisto-bustools and the configuration of alevin-fry that they tested on the `pbmc_10k_v3` data, we do observe some genes as detected. This appears to stem from our use of the most recent release of Seurat (16) (currently version 4.0.0), which modified the default behavior of the `FindMarkers()` function to “prefilter genes and report fold change using base 2, as is commonly done in other differential expression packages, instead of natural log” (https://satijalab.org/seurat/articles/v4_changes.html).

**Fig. 2.**
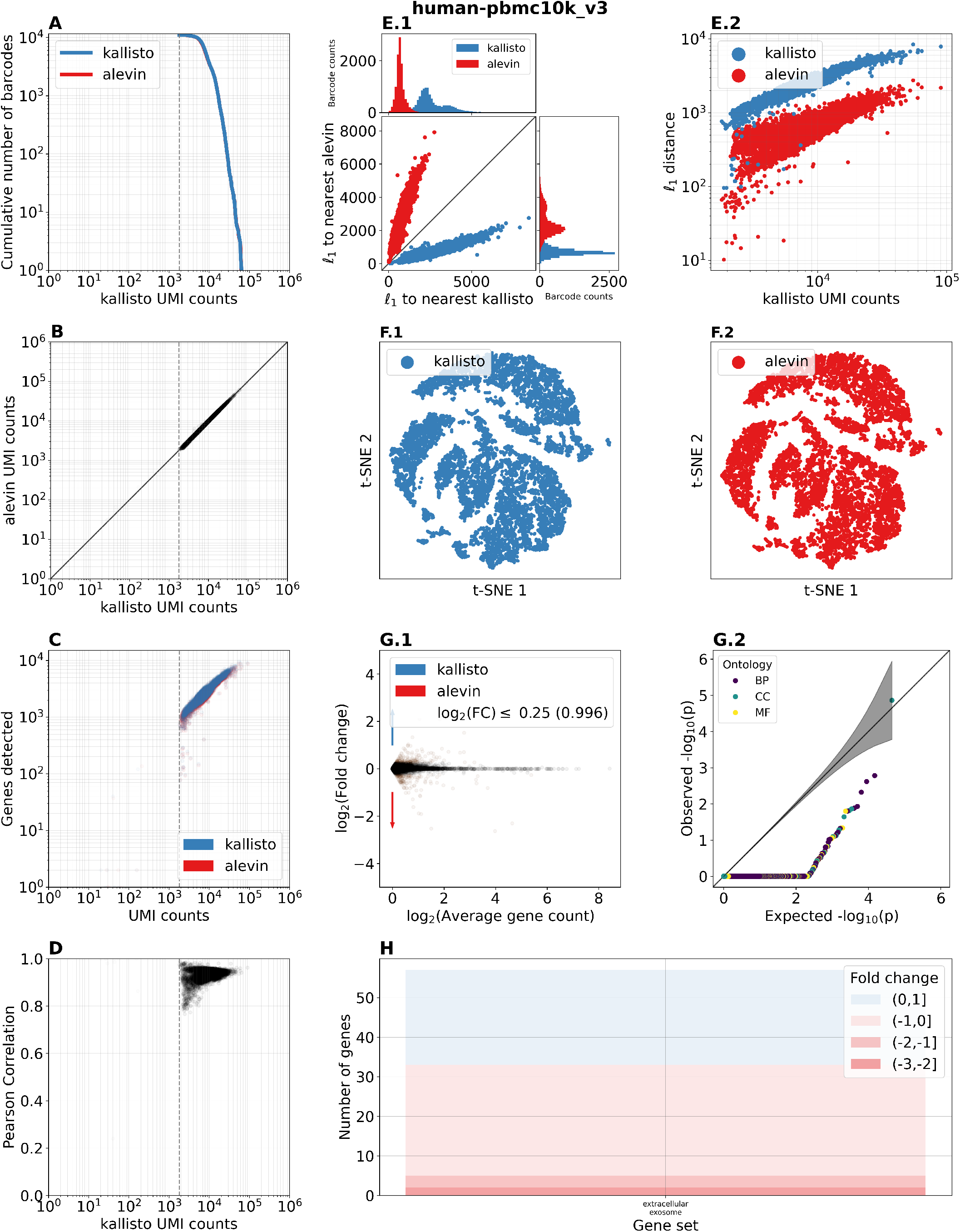
A comparison of the resulting count matrices obtained from alevin-fry and kallisto-bustools, as run in this manuscript, for the `pbmc_10k_v3` dataset. Panels A-H have the same inter-pretation as in Fig. 2 of Booeshaghi and Pachter (1), and compare the count matrices at the gene and cell levels. The plots are generated using the `analysis/scripts/mkplots.py`, `analysis/scripts/mkdata.py` and `analysis/notebooks/run_gsea_bar_full.R` scripts.

For the purposes of keeping this benchmark like-for-like, we have run both tools in configurations where they must generate their own permit list without the input of an external set of valid but unfiltered (*i.e*. possible) barcodes. We choose this configuration for two reasons. First, unlike 10x chromium v2 and v3 experiments, many single-cell technologies supported by both pipelines do not provide an external list of barcodes, and so, this is indicative of the general case where pipelines must provide their own method for generating a permit list of barcodes. Second, since alevin-fry does not yet support unfiltered external permit lists, there is not a way to fairly compare it against a method that can take advantage of this information. We consider it a development priority to support this feature in alevin-fry for single-cell technologies where this information is available. Nonetheless, we tested the effect that requiring a second sort of a (sorted and filtered permit list corrected) BUS file had on the overall runtime, compared to correcting the initial unsorted file with an unfiltered permit list, and processing the unfiltered file in the remainder of the pipeline. To do this, we also ran a configuration of kallisto-bustools where the raw BUS file was first corrected with the external, unfiltered 10x permit list, then the file was sorted, then a permit list was extracted from this sorted file (to allow subsequent filtering of an unfiltered count matrix), and finally, the count step was performed. This process results in an unfiltered matrix which may then be filtered using the generated permit list. Alevin-fry was, on average, ~ 2.15 times as fast as kallisto-bustools under this configuration, rather than ~2.08 as fast; in other words, the runtime costs of the different kallisto-bustools configurations were very similar for these data.

Finally, though we have retained the parallelism settings used in the original benchmark for the purposes of the main results reported in this manuscript, we also evaluated, on one of the larger datasets (`pbmc_10k_v3`), how both tools scale to a higher thread count of 16. In this case, we found that the total runtime for alevin-fry dropped from 19.7 minutes with 10 threads to 15.6 minutes with 16 threads, and the total runtime for kallisto-bustools went from 38.8 minutes with 10 threads to 37.8 minutes with 16 threads. So, in this case, increasing the thread count by 6 lead to a ~ 1.26 times increase in the speed of alevin-fry and a ~1.03 times increase in the speed of kallisto-bustools.

## Conclusions

We find that when alevin-fry is benchmarked in a like-for-like comparison with kallisto-bustools, it is both faster and uses less memory while producing similar results. Of course, in this configuration, alevin-fry (3), unlike the original alevin (13) or other configurations of alevin-fry, is adopting some of the computational simplifications for which Melsted et al. (2) argue, and the similarity of these results is fully expected.

In their manuscript, Booeshaghi and Pachter (1) repeatedly refer to alevin-fry as a “reimplementation” of bustools. This characterization is untrue both in detail and in spirit. The alevin-fry tool has not been designed to reimplement the bustools commands or interface, or specifically to match the implementation of bustools. It has been de-signed as a way to allow the exploration and configuration, in a unified framework, of a variety of different algorithms for single-cell data pre-processing, many of which don’t currently exist in kallisto-bustools. For example, it implements multiple different methods for generating permit lists, and multiple different algorithms for UMI resolution, including some that correct for UMI sequencing errors, resolve multi-gene UMIs by parsimony, probabilistically (which kallisto-bustools subsequently implemented after it was introduced in alevin (13)), or both, as well as the functionality to quantify the uncertainty of probabilistic resolution through bootstrapping. Of course it is the case that, in designing such a tool after the work of Melsted et al. (2) was published and in widespread use, one should learn from the design decisions of that work that proved to be effective and useful. The main such design decisions in this case are first, the separation of the read mapping from the subsequent processing of barcodes and UMIs via intermediate files (as is also done internally by STARsolo), and second, the arrangement of mapping records relevant to a given corrected barcode subsequently so that cells can be processed in an effectively independent manner. The alevin-fry tool adopts these choices described by Melsted et al. (2), as we see no reason to avoid relevant design decisions demonstrated by prior tools, that seem to work, when building new tools. We look forward to discussing these design decisions, as well as some novel design choices we have made, when we publish the alevin-fry preprint.

We have not completed, to our satisfaction, a thorough investigation of the effect of different mapping approaches, permit list generating methods and UMI correction and resolution strategies provided by alevin-fry across a wide range of tagged-end single-cell RNA-seq data and technologies (which have, in general, distinct characteristics compared to both bulk RNA-seq data and full-length single-cell RNA-seq data). Once we have adequately explored this algorithmic parameter space, we plan to publish a full manuscript describing the design and implementation of the alevin-fry pipeline, highlighting where it derives design decisions from kallisto-bustools and other tools, and where it differs, as well as the effect that different configurations have on runtime and memory performance, the raw count matrices and common downstream analyses, and how those effects may vary in different single-cell technologies.

We have described in this manuscript, and demonstrated in the associated code repository and tutorial, how alevin-fry can optionally be configured so as to match the computational complexity of kallisto-bustools as closely as possible. In this like-for-like comparison of these two pipelines, we have shown that, while the estimated gene expressions are similar — at least when assessed in the manner done by Booeshaghi and Pachter (1) — the runtime and memory characteristics are not. Rather, while using the same benchmarking framework as Booeshaghi and Pachter (1), instead of alevin-fry taking ~3 times as long to pre-process data (on average) than kallisto-bustools and using many times as much memory in the worst case (1), we find that alevin-fry is both faster and uses less memory than kallisto-bustools. Specifically, alevin-fry is on average ~ 2.08 times as fast as kallisto-bustools and consumes, on average, only ~0.34 as much peakmemory. According to the formulae used in the jupyter (17) notebooks of Booe-shaghi and Pachter (1) to estimate costs for performing processing on Amazon Web Services compute instances, pre-processing the `pbmc_10k_v3` dataset using the configuration of the alvein-fry pipeline we have tested in this manuscript costs $0.05, which is half of the cost of running the kallisto-bustools pipeline ($0.1). Further-more, if one needs to first build the reference index, the peak runtime memory for kallisto-bustools exceeds 8 GB and so a more expensive instance would be necessary. In that case, building the reference index and processing the `pbmc_10k_v3` would cost $0.23 using kallisto-bustools (while the cost would remain at $0.05 using alevin-fry, even if index construction is included). The cost of the alevin-fry pipeline we have benchmarked in this manuscript is 39 times smaller than what was reported in (1) for this dataset, while the cost of the kallisto-bustools pipeline is twice as large due to the increased memory requirements when using the newer human transcriptome annotation. If one is comfortable with the simplifying assumptions being made, the performance profiles observed in this manuscript provide a compelling case for the use of this configuration of alevin-fry for the rapid and lightweight pre-processing of single-cell RNA-seq data.

Finally, it is important to note that alevin-fry is still undergoing active development and improvement, which is, in part, why no full preprint has yet been published describing the tool and underlying methods and implementation in detail. Of course, one can use the tool today to obtain gene expression counts for single-cell data, but we expect that alevin-fry will continue to advance and expand to offer more capabilities and to be further optimized.

## Footnotes

1 The authors later updated their repository to contain a link to a deposition with the reference data they used, but that information was not available in the original repository commit `d0549e0351c4875428b153c7804f21eed7fa82eb` at the time the preprint was published, and our framework was in place by the time this update was made. This is further described in “Methods.”

2 We do not use the modified version of alevin-fry 0.1.0 that Booeshaghi and Pachter (1) altered to convert encoded barcode identifiers in the `generate-permit-list` step into character strings, but instead the tagged 0.1.0 release with the `rand` crate additionally pinned at 0.7.3 to enable compilation.

3 A subsequent commit included a link to a deposition of the references they used, but our framework was in place and benchmarking underway by the time that commit was made. Further, it is informative to see how even modest changes in the specific reference used can lead to large changes in the memory requirements of a tool.

4 This sketch mode was evaluated in detail in the poster of Sarkar et al. (3), where its scalability was assessed and its mappings were paired with a number of different UMI resolution strategies.

5 The alevin-fry peak memory usage in this dataset happens during the `collate` step, which can easily be made to operate within a strict desired RAM budget; a feature on which we are currently working.

